# Cerebrovascular-balance relationships reveal mechanisms of neuromotor resilience to dual-task cognitive loading

**DOI:** 10.1101/2023.12.19.572430

**Authors:** Jacqueline A. Palmer, Emily M. Hazen, Sandra A. Billinger

**Author notes:** **Correspondence:** Jacqueline A Palmer 420 Delaware St. SE (MMC 388) Minneaopolis, MN 55455 Twitter: @JA_Palmer_.

## Abstract

**BACKGROUND:** Age-related decline in cerebrovascular health precipitates cognitive dysfunction and can be attenuated by habitual physical activity. Cognitive interference in balance control is well-known clinically and reflects compromised neuromotor resilience. However, whether cerebrovascular health affects cognitive-balance dual-tasking with aging is unclear.

**METHODS:** Thirty participants (76±4years) completed clinical balance/cognitive testing under single-task and dual-task conditions. Balance performance was assessed as the total distance traversed during a challenging beam walking task. Cognitive performance was assessed as response time (RT) during a working-memory n-back test. Transcranial Doppler ultrasound was used to measure resting middle cerebral artery blood velocity (MCAv). We tested whether MCAv was associated with single-task and dual-task interference (DTI) in balance and cognitive performance and the effects of age and physical activity level on this relationship.

**RESULTS:** During single-tasking, higher MCAv associated with higher balance function (r=0.40, *p*=0.033), while no relationship was observed for cognitive performance. Dual-tasking strengthened relationships between MCAv and DTI across domains of balance (r=0.442, *p*=0.016) and cognition (r=0.328, *p*=0.089); for cognitive DTI, this effect was driven by individuals ≥75years old (r=0.54, *p*=0.031). Individuals ≥75years exhibited greater cognitive DTI (*p*=0.049), yet achieved similar balance DTI compared to those <75. Regardless of age, participants with higher MCAv demonstrated greater dual-task prioritization of balance control over cognitive performance (r=0.410, *p*=0.030). Physically-active participants with higher MCAv also showed less balance DTI (r=0.621, *p*=0.003), while there was no relationship in under-active/sedentary individuals or within the cognitive domain.

**CONCLUSIONS:** Our results support a key role of cerebrovascular health in neuromotor resilience to cognitive loading, which may emerge earlier in brain aging processes affecting balance control compared to cognition. Cerebrovascular health may support people’s ability to prioritize balance in the face of competing attentional demands, and may mediate the positive effects of physical activity on cortically-mediated balance control.

## 1. Introduction

Cerebrovascular health is emerging as a key neuroprotective factor for neurocognitive function with aging,^1^ yet the role of brain vascular systems in the well-known clinical manifestations of cognitive-motor behavioral interference^2,3^ remain poorly understood. Additionally, relationships between cerebrovascular health and cognitive dysfunction have been well-characterized in the context of disease (e.g. mild cognitive impairment, Alzheimer’s disease, dementia),^4–6^ but remains a nascent area of investigation in older adults with well-preserved cognition (e.g. “super agers”).^7,8^ Additionally, clinical measures of age-related motor dysfunction involving balance and mobility may occur earlier compared to cognitive function, providing early and sensitive clinical biomarkers of brain dysfunction in cognitively-normal older adults.^9–11^ Here, we assessed clinical measures of cerebrovascular health using transcranial Doppler ultrasound and tested its relationship to cognitive-balance dual-task interference in a group of clinically well-characterized, cognitively-normal older adults spanning a wide age range. We further tested whether advanced-age and physical activity level had an effect on the relationship between cerebrovascular health and cognitive-balance dual-task interference.

The emergence of cognitive interference in balance and walking over the course of aging is a prevalent clinical phenomenon^12,13^, e.g. older adults cannot “walk and talk” at the same time.^12^ The ability for cognitive-motor dual-tasking may reflect individual neural capacity^14–16^ that supports neurocognitive and neuromotor resilience imposed by competing attentional demand.^17^ Mobility decline may occur several years before individuals meet clinical diagnosis for mild cognitive impairment (MCI),^9–11^ implicating that motor control may be a more sensitive indicator of underlying neuropathology that precipitates clinical syndrome. Clinical dual-task paradigms can be used to assess cognitive-motor interference, in which the individual is asked to simultaneously perform a cognitive task while balancing/walking and the change in their performance in either or both tasks is measured.^18,19^ Specifically, greater degradation of *balance and walking* performance under cognitive loading (i.e. greater dual-task interference (DTI)) is an early and sensitive indicator of behavioral dysfunction in older adults,^2,3,12,20–23^ and can predict future dementia^24^ and falls.^20,25^ There is also evidence for a cognitive priority “trade-off” with mobility in cognitively-normal older adults who are at high risk for dementia and possess high amyloid-beta deposition in their brains.^18,26^ This poses the question of whether modifiable factors (e.g. vascular health) contribute to the early clinical manifestations of dual-task interference that could be targeted and modified with clinical intervention during preclinical disease stages.

Cerebrovascular health can be quantified clinically and noninvasively as middle cerebral artery blood velocity (MCAv) using transcranial Doppler ultrasound,^27,28^ and has been linked to (single-task) cognitive function in older adults.^29–31^ A decline in MCAv can be observed as early as middle-age^32^ and occurs more slowly in people who are physically-active.^4,33^ Older adults with cerebrovascular disease, possessing lower MCAv and impaired neurovascular coupling, walk at slower gait speeds,^34,35^ implicating a link between cerebrovascular health and mobility in the context of age-related vascular pathology.

In the present study we employ a clinical approach to assess the relationship between cerebrovascular health and cognitive-motor interference in cognitively-normal older adults across a wide age range. We hypothesized that older adults with higher cerebrovascular health, measured using transcranial Doppler ultrasound, would show less cognitive-balance interference, assessed using a dual-task paradigm. We further expected that older adults with higher MCAv would show greater prioritization of balance versus cognitive performance under dual-task conditions, with the strongest effect in individuals in more advanced aging (i.e. ≥75yo). In an exploratory analyses, we tested the effect of physical activity level on the relationship between MCAv and DTI in balance control and cognitive performance.

## 2. Materials and Methods

### 2.1. Participants

Thirty participants (76±4 years, 19 females) from a well-characterized data registry from the University of Kansas Alzheimer’s Disease Research Center (P30AG072973)^3,10–12^ were selected for this study (**Table 1**). Inclusion criteria were (1) age 65-90 years, (2) normal cognition (see below), absence of neurologic or orthopedic disability to prevent independent standing and walking, and (3) English speaking. Exclusion criteria were (1) insulin-dependent diabetes, (2) peripheral neuropathy, (3) active coronary artery disease and congestive heart failure. The University of Kansas Institutional Review Board approved this protocol (IRB#: STUDY 00147888) and all participants provided written informed consent.

**Table 1.** Participant characteristics.

| Table 1. Participant characteristics |  |  |  |  |
| --- | --- | --- | --- | --- |
|  | ALL (n=30) | <75yo (n=13) | ≥75 yo (n=17) | P value |
| Age | 76 ± 4 | 72 ± 1 | 79 ± 4 | *p<0.001 |
| Sex (F/M)<sup>∞</sup> | 19/11 | 11/2 | 8/9 | *p=0.034 |
| BMI | 27 ± 4 | 26 ± 5 | 27 ±4 | p=0.686 |
| Mean arterial pressure (mmHg) | 96 ± 11 | 92 ± 11 | 99 ± 11 | 0.985 |
| Heart rate (bpm) | 91 ± 8 | 90 ± 10 | 99 ± 11 | 0.792 |
| Timed-up-and-go | 7.9±1.4 | 8.0±1.4 | 7.8±1.5 | p=0.796 |
| Gait speed (m/s) | 1.3±0.2 | 1.3±0.2 | 1.3±0.2 | p=0.961 |
| MCAv (cm/s) | 42.7±10.3 | 43.5±11.2 | 42.2±10.0 | p=0.748 |

Values are depicted as mean ± SD. \*p<0.05; ∞self-reported

### 1.1. Clinical screening for cognitive impairment and eligibility

This study specifically aimed to assess clinical cerebrovascular-behavioral relationships in older adults with well-preserved cognition. As such, older adults in the present study underwent a comprehensive neuropsychological test battery and Clinical Dementia Rating (CDR) scale as part of the University of Kansas Alzheimer’s Disease Research Center. A trained clinician and psychometrist performed CDR testing the neuropsychological test battery, respectively. Participants who were rated cognitively-normal (i.e., CDR=0) were eligible for the present study.

### 1.2. Cerebrovascular assessment

Transcranial doppler ultrasound (TCD) was used to assess middle cerebral artery blood velocity (MCAv) during a 5-minute seated rest period. Right MCAv over the temporal window was recorded using a 2-MHz TCD probe (RobotoC2MD, Multigon Industries). The left MCA was used if the right MCA signal was absent. A 5-lead electrocardiogram continuously monitored and recorded heart rhythm. Custom MATLAB software (The Mathworks Inc.) using an analog-to-digital data acquisition unit (NI-USB-6212, National Instruments) acquired MCAv (500 Hz), synchronized across the cardiac cycle.^27,28^ Data were visually inspected and discarded when R-to-R intervals were >5 Hz or changes in peak MCAv exceeded 10 cm/s in a single cardiac cycle. Trials with <85% samples were discarded. Mean MCAv was calculated from the area under the curve for each cardiac cycle.

### 1.3. Behavioral assessments of balance and cognitive performance

A challenging beam walking task was used to assess balance performance.^36–38^ Participants walked across a 16-foot long narrow beam (3.5-inch width) (1-inch height to minimize postural threat).^36^ This dynamic beam walking balance task was selected because it can detect dynamic balance proficiency across older adults and a range of functional sensorimotor abilities,^37^ can detect age-related differences in cognitive-balance dual-task interference,^36^ and may be more sensitive in detecting motor impairments compared to conventional clinical tests.^39^ Older adults who achieved perfect beam scores on the first two trials were progressed to a narrower beam width (1-inch), with a baseline score of 16 feet.

#### Single-task (ST) balance performance

Participants wore a safety belt and were instructed to fold their arms across their chest, fix their gaze at a point on the wall straight ahead at eye level, and walk forward across the beam without stepping off the beam or falling, choosing their own foot placement and walking speed. A trial was stopped when the participant stepped off the beam, walked sideways, or unfolded their arms, and their initial ground foot placement position was marked (**Figure 1A**).

**Figure 1.**
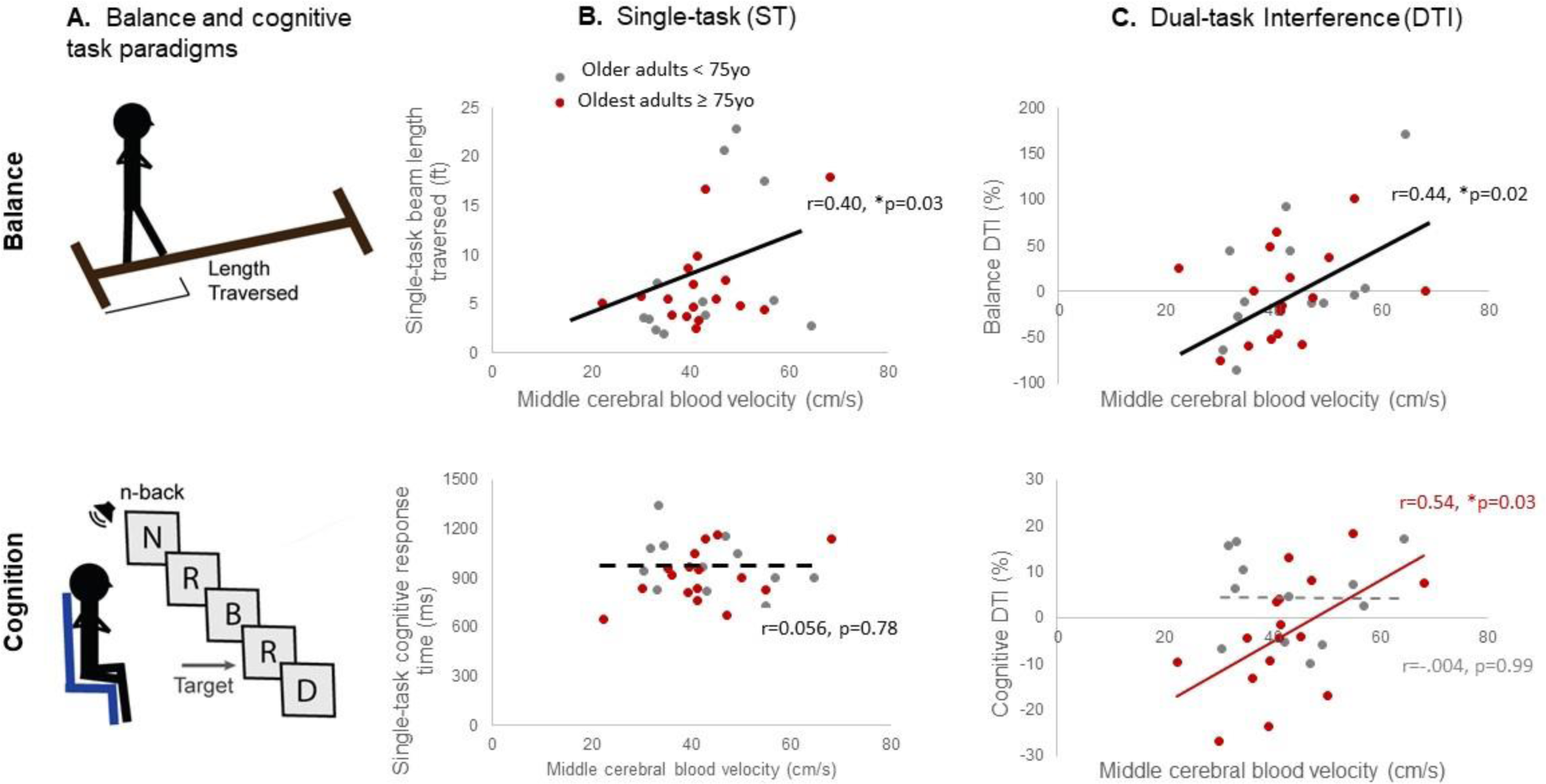
Balance and cognitive performance during single- and dual-task conditions and relationship between middle cerebral artery blood velocity (MCAv). Paradigms for assessing balance and cognitive performance across all older adults are shown **(A)**. During single-task conditions, higher MCAv was associated with higher balance performance regardless of age (r=0.40, p=0.033) while showing no relationship to cognitive performance (**B**). During dual-task conditions, older adults with lower MCAv exhibited a greater decline in beam distance traversed from single- to dual-task conditions (i.e. more negative dual-task interference (DTI)), irrespective of age (r=0.442, p=0.016) **(C,** top). A positive relationship between MCAv and cognitive DTI was driven by older adults ≥ 75 years old (YO) (r=0.54, p=0.031)) **(C,** bottom).

#### Dual-task (DT) balance performance

Following the single-task condition, participants completed a dual-task beam walk during simultaneous performance of a cognitive working-memory task.^36^ The secondary cognitive task required the participant to verbally count backward by 3’s starting at a random integer number between 20 and 100 verbally stated by the experimenter immediately following the cue to begin the beam walk.

The mean beam distance traversed across 2 trials for each single-task and dual-task balance conditions was used to compute balance DTI (below).

#### Single-task (ST) cognitive performance

To enable more precise quantification of cognitive performance, participants completed a cognitive working-memory task using the auditory n-back (2-n) test delivered through E-Prime Software (Pittsburgh, PA). The n-back test is widely validated in older adult populations.^10^ Participants were asked to recall a repeated letter from 2 letters prior in a 50-letter sequence delivered at 0.5Hz by clicking a button on a hand-held clicker as accurately and as quickly as possible (**Figure 1A**).

#### Dual-task (DT) cognitive performance

Following the single-task condition, participants completed the cognitive n-back test while simultaneously standing on an unstable balance board (Fluidstance Level Balance Board) (Santa Barbara, CA). The level of balance challenge was individually adjusted with the addition or removal of a convex board cap based on the participant’s balance ability, determined by a licensed physical therapist. Participants were required to fold their arms across their chest and affix their gaze at an eye-level point on the wall while holding the remote clicker.

The mean response time across all accurate trials in each the single- and dual-task conditions was used to compute *dual-task interference (DTI)* as:

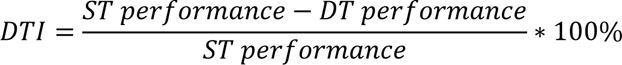

For balance DTI, the value was multiplied by (-1), such that a negative DTI value for each balance and cognition indicates worsening of performance between the single- and dual-task conditions.^19^

Additionally, *prioritization* of balance versus cognition was computed as:

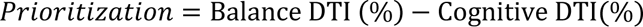

A positive value indicates prioritization of balance over cognitive performance (i.e. less DTI in balance relative to cognitive performance).

#### Physical activity level

As a secondary outcome, physical activity level was screened using the Rapid Assessment of Physical Activity (RAPA) self-report questionnaire, and participants were classified as physically-active or under-active/sedentary.^41^ The RAPA shows good or better sensitivity and predictive value compared to other self-assessment tools for physical activity in older adult populations.^41^

### 1.4. Statistical analyses

We confirmed normality and heterogeneity of variance using Kolmogorov-Smirnov and Levene’s tests, respectively. We used Pearson product-moment correlation coefficients to test relationships between MCAv and each balance and cognitive performance during single-task and DTI. Given the nonlinear effect of advanced aging (≥75yo),^42–44^ we then used two-way multiple linear regression (MLR) analyses (factors: age, MCAv, age-by-MCAv) to test whether cerebrovascular-behavioral relationships differed as a function of age in dichotomized subgroups (≥75yo and <75yo).

We then tested the effects of age on prioritization of balance versus cognitive performance; we used 1) independent t-tests to compare balance and cognitive DTI between age groups and 2) a chi-square test to examine the relation between age and prioritization of balance versus cognition. We tested for an interactive age-by-MCAv effect on prioritization using MLR analysis. In an exploratory analyses, we tested whether relationships between MCAv and DTI were different in physically-active compared to under-active/sedentary older adults across domains of balance and cognition. All analyses were performed using SPSS version 29 with an *a priori* level of significance set to 0.05.

## RESULTS

Two participants had no right side MCA signal, so the left MCA was used. One participant did not have an MCA signal in either L or R side; their data were excluded from all analyses involving MCAv. One participant declined to perform the dual-task n-back test; their data were excluded from all analyses involving cognitive performance and DTI. Participants in the subgroup ≥75 years differed in age by a mean of 7 years compared to those in the younger subgroup and had a greater proportion of male participants (*p*=0.034). In contrast to expected functional differences between these age groups, the subgroup of older adults ≥75 years showed no difference in standard clinical measures of mobility function on the timed-up-and-go test (*p*=0.796), walking speed *(p*=0.961), and no difference in MCAv (*p*=0.748) (see **Table 1**).

### Relationship between cerebral blood flow and behavior across domains of cognition and balance control and effect of dual-task interference

During single-task performance, there was a positive association between MCAv and balance performance for the entire group (r=0.40, *p*=0.033), which was unaffected by participant age (model, *p*=0.207; interaction, *p*=0.917). Conversely, there was no discernible correlation between MCAv and single-task cognitive performance for the whole group (r=0.056, *p*=0.778), and the effect of age was negligible (model, *p*=0.119; interaction, p=0.037) (**Figure 1B**).

Under dual-task interference, cerebrovascular-behavioral relationships were strengthened across balance and cognitive behavior. Within the balance domain, older adults with lower MCAv exhibited a greater decline in beam distance traversed from single-to dual-task conditions (r=0.442, *p*=0.016), irrespective of age (model, *p*=0.069; interaction, *p*=0.284). In the cognitive domain, dual-tasking revealed a trend towards a positive relationship between MCAv and cognitive dual-task interference behavior for the entire group (r=0.328, *p*=0.089). This effect was driven by older adults aged 75 and above (r=0.54, *p*=0.031) but was absent in individuals under 75 years old (r=-0.004, *p*=0.991). Notably, when assessing differences in relationships between age groups, there was a trend towards an age-by-MCAv interaction (model *p*=0.030; interaction, *p*=0.100) (**Figure 1C**).

### Effects of age and cerebral blood flow on the prioritization of balance vs cognition

When comparing dual-task interference (DTI) between age subgroups, the oldest subgroup exhibited greater decline in cognitive performance during dual-tasking (i.e., greater cognitive DTI) (*p*=0.049), while showing no significant difference in balance DTI (*p*=0.709) (**Figure 2A**). In evaluating the prioritization of balance versus cognition during dual-tasking, the relationship between age subgroup (≥ or < 75 years old) and the priority of balance over cognition was not statistically significant [*X^2^* (1, N=29 = 0.003, p = 0.628)]. Individuals aged ≥75 years were no less likely to prioritize balance over cognition during dual-tasking compared to individuals approximately 7 years younger (**Figure 2B**).

**Figure 2.**
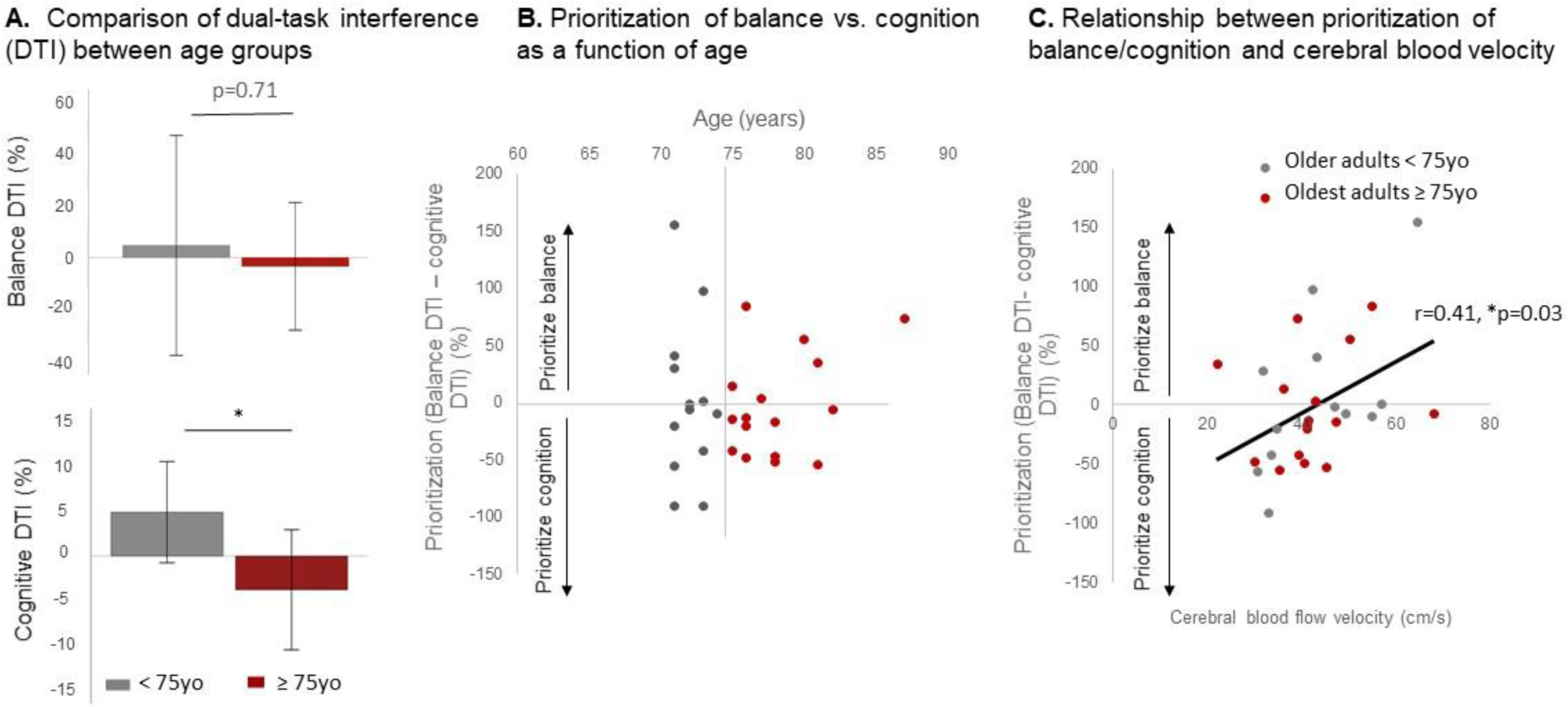
Prioritization of balance versus cognitive performance and relationships to middle cerebral artery blood velocity (MCAv). Older adults ≥ 75 years old (yo) showed greater cognitive DTI compared to those < 75yo (\**p*=0.049), but exhibited no significant difference in balance DTI (*p*=0.709) **(A).** There was no relationship between age subgroup (≥ or < 75 years old) and the priority of balance over cognition [*X^2^* (1, N=29 = 0.003, p = 0.628)] **(B)**. Higher MCAv was associated with increased prioritization of balance control over cognition across all older adults, regardless of age(r=0.410, p=0.030) **(C).**

Further, it was observed that older adults with higher MCAv demonstrated an increased prioritization of balance control over cognition (whole group r=0.410, *p*=0.030). This effect persisted regardless of age (model: *F*_3,27_=2.564, *p*=0.078, *R^2^*=0.243, adjusted *R^2^*=0.148; interaction: t= -1.473, p=0.154) (**Figure 2C**).

### Effect of physical activity on the relationship between cerebral blood flow and dual-task interference

When examining the influence of individual physical activity levels on relationships between MCAv and dual-task interference (DTI), distinct effects were observed between balance and cognitive performance. In the domain of balance, there was a significant interactive effect of physical activity with MCAv on balance DTI (model: *F*_3,28_=4.18, *p*=0.016, *R^2^*=0.334, adjusted *R^2^*=0.254; interaction: t= -2.26, *p*=0.033). Specifically, physically-active older adults with higher MCAv exhibited less decline in balance control during dual-tasking (r=0.621, p=0.003), while sedentary older adults showed no discernible relationship (r=0.101, *p*=0.812) (**Figure 3A**).

**Figure 3.**
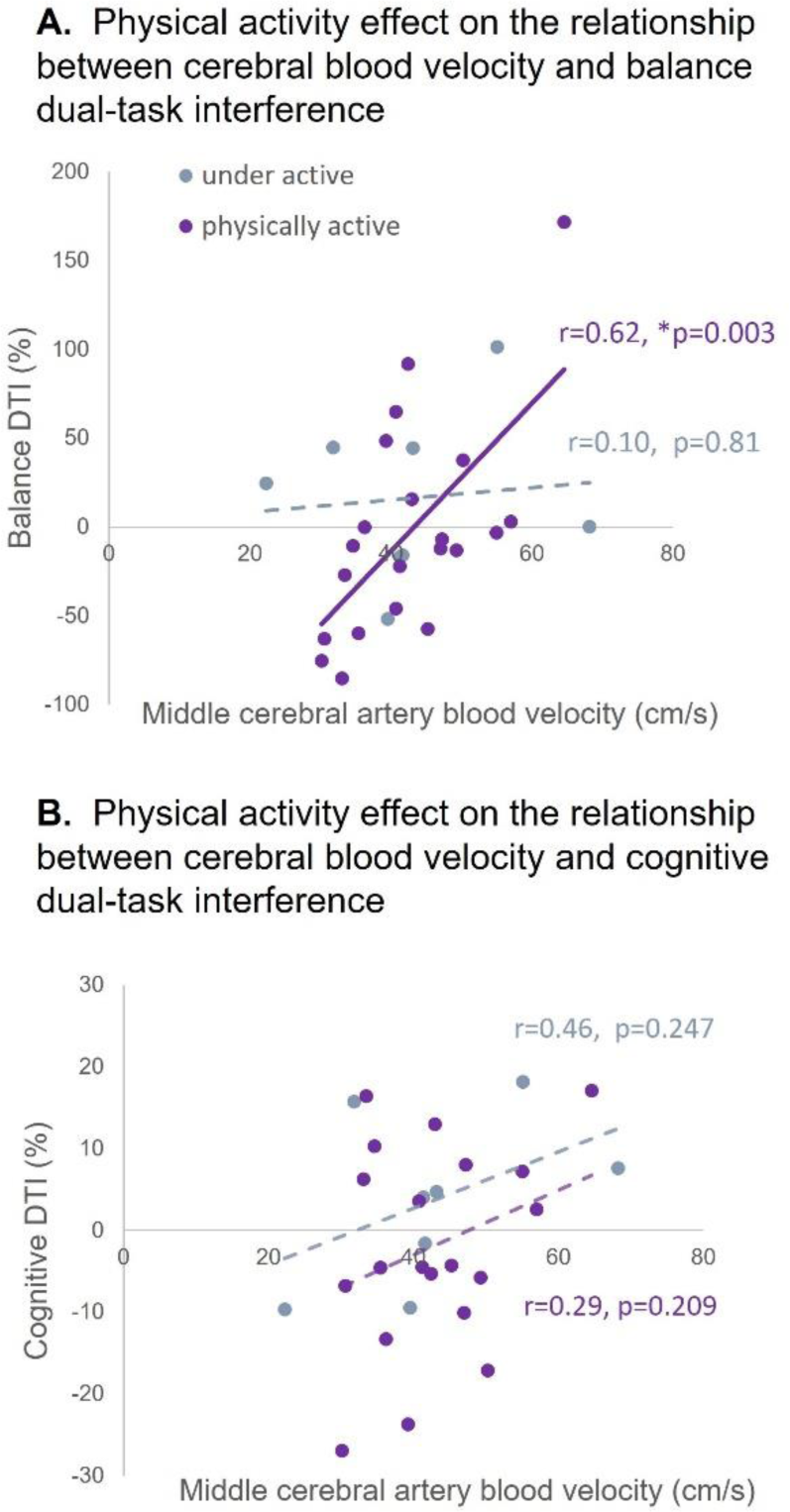
**Relationship between middle cerebral artery blood velocity (MCAv) and dual-task interference (DTI) in physically-active and under-active/sedentary older adults** in each domain of balance **(A)** and cognitive performance **(B)**. There was a significant interactive effect of physical activity with MCAv on balance DTI (*F*_3,28_=4.18, *p*=0.016, interaction: t= -2.26, *p*=0.033) in which physically-active older adults with higher MCAv exhibited less balance DTI (r=0.621, *p*=0.003), while under-active/sedentary older adults showed no relationship. Physical activity level did not influence the relationship between MCAv and cognitive DTI, nor was there a significant relationship between MCAv and cognitive DTI in physically-active or under-active/sedentary older adults without controlling for age.

Conversely, in the domain of cognition, physical activity level did not influence the relationship between MCAv and cognitive DTI (model: *F*_3,27_=1.45, *p*=0.254, *R^2^*=0.152, adjusted *R^2^*=0.047; interaction: t= 0.115, *p*=0.909). There was no significant relationship between MCAv and cognitive DTI in physically-active (r=0.294, *p*=0.209) or sedentary/underactive older adults (r=0.464, *p*=0.247) without controlling for age (**Figure 3B**).

## DISCUSSION

Findings of the present study provide initial evidence that clinically-meaningful measures of cognitive-motor interference and balance prioritization during dual-tasking are closely related to cerebrovascular health. Further, results show that cerebrovascular health interacts with individual physical activity level in older adults who are resistant to cognitive decline. The strengthening of relationships between MCAv and higher balance and cognitive function under dual-task conditions implicate an increased role of cerebrovascular health for behaviors that necessitate greater cortical resources. Additionally, the increased role of cerebrovascular health may occur earlier for aging brain processes affecting balance control compared to cognition, as the link between MCAv and cognitive performance emerged only in the oldest individuals ≥75years. Notably, individuals with higher MCAv demonstrated a greater prioritization of balance over cognitive performance regardless of age, suggesting that cerebrovascular health may support cortically-mediated balance control in the presence of competing attentional demands, potentially enabling older adults over the age of 75 to achieve comparable clinical dual-task balance control to individuals approximately 7 years younger. Lastly, our findings offer preliminary evidence for cerebrovascular health as a mediator for the neuroprotective effects of physical activity on clinical dual-task balance function in cognitively-normal older adults. Our findings elucidate the integral role of cerebrovascular health in supporting brain function for interactive cognitive and motor behaviors and extends previous research in disease states to neurotypical older adults who are resistant cognitive decline, even in the presence of advanced age. Importantly, these results underscore cerebrovascular health as a promising target for interventions aimed at maximizing neurocognitive and neuromotor resilience linked to age-related disease.

### Challenging balance control reveals cerebrovascular-behavioral relationships in older adults resistant to cognitive decline

Our study provides evidence that challenging balance control serves as a sensitive clinical probe to investigate cerebrovascular-behavioral relationships in cognitively-normal older adults, potentially offering early mechanistic insight into neuromotor resilience with aging. Existing evidence shows that brain resilience, the ability to cope with challenge, declines with age and influences the susceptibility to dementia.^1^ Simultaneous cognitive loading during balance control may probe neurocognitive and neuromotor resilience to physiologic challenge.^17^ While the precise neural mechanisms underpinning the interplay between cognition and motor control is unclear, one theory poses that dual-task performance is limited by an individuals’ neural capacity that ultimately limits behavioral task complexity.^14–16^ Building upon prior studies in single-task conditions,^34,35,45^ our results support the notion that brain resilience and functional capacity are challenged by cognitive-balance dual-tsking, and that higher cerebrovascular health may promote greater neurocognitive and neuromotor resilience to this physiologic challenge in cognitively-normal older adults

We found that challenging balance control unveiled cerebrovascular-behavior relationships across a wide age range for balance performance, which were masked during single-task cognitive assessments (see **Figure 1**), and are consistent with motor impairment that precipitates clinical cognitive dysfunction in this preclinical population.^9–11^ Notably, the emergence of cerebrovascular-cognitive relationships under dual-task conditions was exclusive to individuals over 75 years old (see **Figure 1C**). These findings suggest that cognitively-normal older adults exhibit less resilience to the unique neuromotor challenge presented by balance control and cognitive-balance dual-task assessments. From a clinical perspective, these results support the use of balance testing to expose early biomarkers of brain dysfunction and factors for brain resiliency (e.g. higher cerebrovascular health) in coping with physiologic challenge.

### Greater cerebrovascular health may support older adults’ ability to prioritize balance over cognition during dual tasking

In the face of competing attentional demand, the present findings suggest that greater cerebrovascular health may support older adults’ ability to prioritize balance over cognition, an effect that persists regardless of age. As expected based on previous reports of age-related decline in cognitive performance,^7^ individuals ≥75years old demonstrated compromised cognitive DTI. Yet, surprisingly, these individuals showed no difference in balance dual-task interference (DTI) between age subgroups (**Figure 2A&B**). When examining the relationship between cerebrovascular health and cognitive-balance prioritization, we found that older adults with higher MCAv showed a greater prioritization of balance over cognition during dual-tasking, irrespective of age (**Figure 2C**). This is an interesting finding, given the differential effect of age on the relationship between MCAv and balance DTI compared to cognition, as depicted in **Figure 1**, implicating that greater cerebrovascular health may support effective cortical compensation for balance control with aging.^12,13^ This cortical strategy may be unmasked in the context of competing attentional demands between motor and cognitive behaviors, revealing greater cognitive performance compromise in the oldest subgroup.

Alterations in cognitive-balance prioritization is observed in age-related disease such as Parkinson’s disease, MCI, and dementia, in which dual-task costs to gait speed exceed those to cognition.^24,46^ Longitudinal examinations post-stroke show a transition from mutual cognitive and motor dual-task interference to more exaggerated cognitive interference in motor behavior.^47^ Higher genetic risk for Alzheimer’s disease^18^ and greater structural brain pathology (i.e., amyloid-beta)^26^ is also associated with a preference for cognition over mobility in older adults. These findings serve as a foundation for clinical trials to test whether clinical intervention-induced improvements in cerebrovascular health (e.g. through increased physical activity)^4,33^ may prevent a maladaptive tradeoff of cognitive performance over balance control and associated neuropathology.

### Physically-active older adults show an increased role of cerebrovascular health in neuromotor resilience to cognitive loading

These findings provide evidence that cerebrovascular health may mediate the neuroprotective effects of physical activity on brain function uniquely for dual-task balance control during preclinical aging processes. The motor specificity and the complex, multifaceted effects of physical activity^48^ may explain the unique interactive effects of physical activity with MCAv for balance DTI in older adults that did not extend to the domain of dual-task cognitive performance (**Figure 3**). Future clinical trials employing exercise interventions may consider the utilization of clinical transcranial ultrasound to assess cerebrovascular health and it’s relationship to exercise-induced behavioral and neural plasticity.

### Strengths and limitations

Subjecting participants to a comprehensive cognitive battery enabled us to confidently capture cognitively-normal older adults in advanced aging, who may be underrepresented in cognitively-normal cohorts given the pivotal role age plays in cognitive decline.^49^ Nevertheless, the modest sample size and cross-sectional nature underscores the need for larger investigations with longitudinal measures. Differences in self-reported sex between age groups in the present study could have influenced cerebrovascular-behavioral relationships^30^ and warrants further investigation. While transcranial ultrasound offers clinical applicability, it offers limited regional specificity compared to MR-based techniques.^50^ The focus on a single cognitive domain across balance dual-task conditions restricts interpretation to working memory contexts. Variations may arise when applying this paradigm to other cognitive domains (e.g., executive function).^29^ The reliance on self-reported physical activity necessitates confirmation using objective measures, potentially facilitated by wearable technology. The generalizability and reproducibility of our findings are constrained by the underrepresentation of non-white races as a result of recruitment in these segments of community-dwelling older adults, emphasizing the necessity for increased outreach efforts.

## CONCLUSION

The present findings identify cerebrovascular health as a mechanism supporting neuromotor resilience to cognitive loading in older adults who are resistant to cognitive decline. Our findings from a model of healthy aging offer the potential to create a paradigm shift in the clinical and scientific framework of balance control with aging, suggesting that physical activity interventions targeting increased cerebral blood flow may offer an effective method to delay or prevent early clinical manifestations of cognitive interference in balance control.

## Author contributions

JP conceived of the present research question and designed this study. JP and SB obtained funding to support this study. JP and EH collected and analyzed the data. SB, EH, and the KU Alzheimer’s Disease Research Center contributed to recruitment. JP constructed the figures and the first draft of this manuscript. SB contributed the analysis tools and expertise for interpreting cerebrovascular data. All authors discussed the results and contributed to the final manuscript.

## REFERENCES

1. Iturria-Medina Y, Sotero RC, Toussaint PJ, Mateos-Pérez JM, Evans AC. Early role of vascular dysregulation on late-onset Alzheimer’s disease based on multifactorial data-driven analysis. Nature Communications. 2016;7(1):11934. doi:10.1038/ncomms11934

2. Woollacott M, Shumway-Cook A. Attention and the control of posture and gait: a review of an emerging area of research. Gait & Posture. 2002;16(1):1–14. doi:10.1016/S0966-6362(01)00156-4

3. Brown LA, Shumway-cook A, Woollacott MH. Attentional Demands and Postural Recovery : The Effects ofAging. Journal of Gerontology. 1999;54(4):165–171.

4. Tarumi T, Zhang R. Cerebral blood flow in normal aging adults: cardiovascular determinants, clinical implications, and aerobic fitness. Journal of Neurochemistry. 2018;144(5):595–608. 10.1111/jnc.14234

5. Ogoh S. Relationship between cognitive function and regulation of cerebral blood flow. J Physiol Sci. 2017;67(3):345–351. doi:10.1007/s12576-017-0525-0

6. Stefanidis KB, Isbel B, Klein T, Lagopoulos J, Askew CD, Summers MJ. Reduced cerebral pressure-flow responses are associated with electrophysiological markers of attention in healthy older adults. J Clin Neurosci. 2020;81:167–172. doi:10.1016/j.jocn.2020.09.034

7. Gefen T, Shaw E, Whitney K, et al. Longitudinal Neuropsychological Performance of Cognitive SuperAgers. J Am Geriatr Soc. 2014;62(8):1598–1600. doi:10.1111/jgs.12967

8. Cook AH, Sridhar J, Ohm D, et al. Rates of Cortical Atrophy in Adults 80 Years and Older With Superior vs Average Episodic Memory. JAMA. 2017;317(13):1373–1375. doi:10.1001/jama.2017.0627

9. Hoogendijk EO, Rijnhart JJM, Skoog J, et al. Gait speed as predictor of transition into cognitive impairment: Findings from three longitudinal studies on aging. Experimental Gerontology. 2020;129:110783. doi:10.1016/j.exger.2019.110783

10. Rosano C, Snitz BE. Predicting dementia from decline in gait speed: Are we there yet? J Am Geriatr Soc. 2018;66(9):1659–1660. doi:10.1111/jgs.15368

11. Quan M, Xun P, Chen C, et al. Walking Pace and the Risk of Cognitive Decline and Dementia in Elderly Populations: A Meta-analysis of Prospective Cohort Studies. The Journals of Gerontology: Series A. 2017;72(2):266–270. doi:10.1093/gerona/glw121

12. Lundin-Olsson L, Nyberg L, Gustafson Y. “Stops walking when talking” as a predictor of falls in elderly people. Lancet. 1997;349(9052):617. doi:10.1016/S0140-6736(97)24009-2

13. Duckrow R. Stance perturbation-evoked potentials in old people with poor gait and balance. Clinical Neurophysiology. 1999;110(12):2026–2032. doi:10.1016/S1388-2457(99)00195-9

14. Tombu M, Jolicoeur P. Testing the predictions of the central capacity sharing model. J Exp Psychol Hum Percept Perform. 2005;31(4):790–802. doi:10.1037/0096-1523.31.4.790

15. Tombu M, Jolicoeur P. All-or-none bottleneck versus capacity sharing accounts of the psychological refractory period phenomenon. Psychol Res. 2002;66(4):274–286. doi:10.1007/s00426-002-0101-x

16. Tombu M, Jolicoeur P. A central capacity sharing model of dual-task performance. J Exp Psychol Hum Percept Perform. 2003;29(1):3–18. doi:10.1037//0096-1523.29.1.3

17. Montine TJ, Cholerton BA, Corrada MM, et al. Concepts for brain aging: resistance, resilience, reserve, and compensation. Alzheimer’s Research & Therapy. 2019;11(1):22. doi:10.1186/s13195-019-0479-y

18. Whitson HE, Potter GG, Feld JA, et al. Dual-Task Gait and Alzheimer’s Disease Genetic Risk in Cognitively Normal Adults: A Pilot Study. J Alzheimers Dis. 2018;64(4):1137–1148. doi:10.3233/JAD-180016

19. Plummer P, Eskes G. Measuring treatment effects on dual-task performance: a framework for research and clinical practice. Front Hum Neurosci. 2015;9. doi:10.3389/fnhum.2015.00225

20. Montero-Odasso M, Verghese J, Beauchet O, Hausdorff JM. Gait and cognition: a complementary approach to understanding brain function and the risk of falling. J Am Geriatr Soc. 2012;60(11):2127–2136. doi:10.1111/j.1532-5415.2012.04209.x

21. Leone C, Feys P, Moumdjian L, D’Amico E, Zappia M, Patti F. Cognitive-motor dual-task interference: A systematic review of neural correlates. Neuroscience & Biobehavioral Reviews. 2017;75:348–360. doi:10.1016/j.neubiorev.2017.01.010

22. Morris R, Lord S, Bunce J, Burn D, Rochester L. Gait and cognition: Mapping the global and discrete relationships in ageing and neurodegenerative disease. Neuroscience and Biobehavioral Reviews. 2016;64:326–345. doi:10.1016/j.neubiorev.2016.02.012

23. Rankin JK, Woollacott MH, Shumway-cook A, Brown LA. Cognitive Influence on Postural Stability : A Neuromuscular Analysis in Young and Older Adults. Journal of Gerontology. 2000;55(3):112–119.

24. Montero-Odasso MM, Sarquis-Adamson Y, Speechley M, et al. Association of Dual-Task Gait With Incident Dementia in Mild Cognitive Impairment: Results From the Gait and Brain Study. JAMA Neurol. 2017;74(7):857–865. doi:10.1001/jamaneurol.2017.0643

25. Shumway-Cook A, Woollacott M, Kerns KA, Baldwin M. The Effects of Two Types of Cognitive Tasks on Postural Stability in Older Adults With and Without a History of Falls. The Journals of Gerontology Series A: Biological Sciences and Medical Sciences. 1997;52A(4):M232-M240. doi:10.1093/gerona/52A.4.M232

26. Nadkarni NK, Lopez OL, Perera S, et al. Cerebral Amyloid Deposition and Dual-Tasking in Cognitively Normal, Mobility Unimpaired Older Adults. The Journals of Gerontology Series A: Biological Sciences and Medical Sciences. 2017;72(3):431. doi:10.1093/gerona/glw211

27. Billinger SA, Craig JC, Kwapiszeski SJ, et al. Dynamics of middle cerebral artery blood flow velocity during moderate-intensity exercise. J Appl Physiol (1985). 2017;122(5):1125-1133. doi:10.1152/japplphysiol.00995.2016

28. Ward JL, Craig JC, Liu Y, et al. Effect of healthy aging and sex on middle cerebral artery blood velocity dynamics during moderate-intensity exercise. Am J Physiol Heart Circ Physiol. 2018;315(3):H492–H501. doi:10.1152/ajpheart.00129.2018

29. Palmer JA, Kaufman CS, Vidoni ED, Honea RA, Burns JM, Billinger SA. Cerebrovascular response to exercise interacts with individual genotype and amyloid-beta deposition to influence response inhibition with aging. Neurobiology of Aging. Published online March 4, 2022. doi:10.1016/j.neurobiolaging.2022.02.014

30. Palmer JA, Kaufman CS, Vidoni ED, Honea RA, Burns JM, Billinger SA. Sex Differences in Resilience and Resistance to Brain Pathology and Dysfunction Moderated by Cerebrovascular Response to Exercise and Genetic Risk for Alzheimer’s Disease. J Alzheimers Dis. Published online September 19, 2022. doi:10.3233/JAD-220359

31. Alwatban MR, Aaron SE, Kaufman CS, et al. Effects of age and sex on middle cerebral artery blood velocity and flow pulsatility index across the adult lifespan. J Appl Physiol (1985). 2021;130(6):1675-1683. doi:10.1152/japplphysiol.00926.2020

32. Xing CY, Tarumi T, Liu J, et al. Distribution of cardiac output to the brain across the adult lifespan. J Cereb Blood Flow Metab. 2017;37(8):2848–2856. doi:10.1177/0271678X16676826

33. Thomas BP, Yezhuvath US, Tseng BY, et al. Life-long aerobic exercise preserved baseline cerebral blood flow but reduced vascular reactivity to CO2. Journal of magnetic resonance imaging: JMRI. 2013;38(5):1177–1183. doi:10.1002/jmri.24090

34. Sorond FA, Galica A, Serrador JM, et al. Cerebrovascular hemodynamics, gait, and falls in an elderly population: MOBILIZE Boston Study. Neurology. 2010;74(20):1627–1633. doi:10.1212/WNL.0b013e3181df0982

35. Sorond FA, Kiely DK, Galica A, et al. Neurovascular coupling is impaired in slow walkers: The MOBILIZE Boston Study. Annals of Neurology. 2011;70(2):213–220. doi:10.1002/ana.22433

36. Uematsu A, Tsuchiya K, Suzuki S, Hortobágyi T. Cognitive dual-tasking augments age-differences in dynamic balance quantified by beam walking distance: A pilot study. Exp Gerontol. 2018;114:27–31. doi:10.1016/j.exger.2018.10.016

37. Sawers A, Ting LH. Gait & Posture Beam walking can detect differences in walking balance proficiency across a range of sensorimotor abilities. Gait & Posture. 2015;41(2):619–623. doi:10.1016/j.gaitpost.2015.01.007

38. Kemp GJ, Birrell F, Clegg PD, et al. Developing a toolkit for the assessment and monitoring of musculoskeletal ageing. Age Ageing. 2018;47(suppl_4):iv1-iv19. doi:10.1093/ageing/afy143

39. Hortobágyi T, Uematsu A, Sanders L, et al. Beam Walking to Assess Dynamic Balance in Health and Disease: A Protocol for the “BEAM” Multicenter Observational Study. Gerontology. 2019;65(4):332–339. doi:10.1159/000493360

40. Bopp KL, Verhaeghen P. Aging and n-Back Performance: A Meta-Analysis. J Gerontol B Psychol Sci Soc Sci. 2020;75(2):229–240. doi:10.1093/geronb/gby024

41. Topolski TD, LoGerfo J, Patrick DL, Williams B, Walwick J, Patrick MB. The Rapid Assessment of Physical Activity (RAPA) among older adults. Prev Chronic Dis. 2006;3(4):A118.

42. Osoba MY, Rao AK, Agrawal SK, Lalwani AK. Balance and gait in the elderly: A contemporary review. Laryngoscope Investig Otolaryngol. 2019;4(1):143–153. doi:10.1002/lio2.252

43. Alshammari SA, Alhassan AM, Aldawsari MA, et al. Falls among elderly and its relation with their health problems and surrounding environmental factors in Riyadh. J Family Community Med. 2018;25(1):29–34. doi:10.4103/jfcm.JFCM_48_17

44. Freedman VA, Kasper JD, Spillman BC, et al. Behavioral Adaptation and Late-Life Disability: A New Spectrum for Assessing Public Health Impacts. Am J Public Health. 2014;104(2):e88–e94. doi:10.2105/AJPH.2013.301687

45. Jor’dan AJ, Poole VN, Iloputaife I, et al. Executive Network Activation is Linked to Walking Speed in Older Adults: Functional MRI and TCD Ultrasound Evidence From the MOBILIZE Boston Study. J Gerontol A Biol Sci Med Sci. 2017;72(12):1669–1675. doi:10.1093/gerona/glx063

46. Al-Yahya E, Dawes H, Smith L, Dennis A, Howells K, Cockburn J. Cognitive motor interference while walking: a systematic review and meta-analysis. Neurosci Biobehav Rev. 2011;35(3):715–728. doi:10.1016/j.neubiorev.2010.08.008

47. Liu YC, Yang YR, Tsai YA, Wang RY. Cognitive and motor dual task gait training improve dual task gait performance after stroke - A randomized controlled pilot trial. Sci Rep. 2017;7(1):4070. doi:10.1038/s41598-017-04165-y

48. Eggenberger P, Wolf M, Schumann M, de Bruin ED. Exergame and Balance Training Modulate Prefrontal Brain Activity during Walking and Enhance Executive Function in Older Adults. Front Aging Neurosci. 2016;8:66. doi:10.3389/fnagi.2016.00066

49. Rogalski EJ. Don’t forget—Age is a relevant variable in defining SuperAgers. Alzheimers Dement (Amst). 2019;11:560–561. doi:10.1016/j.dadm.2019.05.008

50. Palmer JA, Morris JK, Billinger SA, et al. Hippocampal blood flow rapidly and preferentially increases after a bout of moderate-intensity exercise in older adults with poor cerebrovascular health. Cereb Cortex. Published online October 18, 2022:bhac418. doi:10.1093/cercor/bhac418

